# demres: An R package to study time-varying demographic resilience

**DOI:** 10.1101/2024.12.10.627540

**Authors:** Julie Louvrier, Alexandre Courtiol, Ella W. White, Adam T. Clark, Sarah Benhaiem, Viktoriia Radchuk

## Abstract

1. Quantifying the resilience of populations to disturbances is essential to assess how threatened populations are. Demographic resilience has been defined as the ability of populations to resist and recover from alterations in their demographic structures. Resilience metrics are typically obtained by applying transient analyses to matrix population models. Until now, the questions of how demographic resilience can change over time, how to quantify such temporal variation and when it is important to account for it have not yet been studied. However, since demographic rates fluctuate over time, it is essential to evaluate whether and under what conditions the assumption of time-constant demographic resilience remains appropriate.
2. In this study we introduce demres - an R package that offers functions to quantify time- varying and time-constant demographic resilience metrics, including time to convergence, damping ratio, inertia, maximum amplification, maximum attenuation and reactivity. The package also allows the comparison of these two approaches, both visually and by means of distance metrics.
3. We use a case study to illustrate the versatility of demres and demonstrate how this tool can be readily used by conservation biologists, managers and others to assess the time- varying resilience of wildlife populations using long-term demographic data. Our framework facilitates standardised comparisons of demographic resilience metrics, for example by means of comparative studies across populations and species.

## Introduction

Quantifying resilience is essential to predict the fate of populations living in environments that are undergoing change. Capdevila et al. (2020) defined demographic resilience as the ability of populations to resist and recover from alterations in their demographic structures. By measuring demographic resilience we can assess immediate, short-term, response of populations to disturbances, which proved useful when assessing for example the influence of environmental stochasticity on populations (Gilbert et al., 2020). Demographic resilience metrics are quantified by applying matrix population models (MPMs, Caswell, 2006) analyse the short-term (i.e. transient) response of populations to disturbances (Stott et al., 2011, 2012). As input, MPMs use vital rates that summarise the survival and fecundity of age or stage classes of the population. The common approach to quantify demographic resilience when vital rates have been collected over a certain period of time is to average them over that period and to use a so-called time-averaged matrix (Capdevila et al., 2020, 2021). This approach is generally common practice in classical demographic analyses of MPMs that focus on long-term (i.e. asymptotic) population dynamics (Caswell, 2006). In the case of transient analysis that focuses on short-term dynamics, the averaging of vital rates over time implicitly assumes that they do not vary strongly enough in time to warrant quantification of temporally-varying demographic resilience. However, it is unclear whether this assumption is justified. As vital rates are reported to vary strongly over time across species and study systems (Bailey et al., 2024; Jenouvrier et al., 2022; Jonzén et al., 2010; Marescot et al., 2018) so are demographic resilience metrics likely to vary over time too.

Long-term studies of populations in the wild show that vital rates can vary strongly over time (Bailey et al., 2024; Jenouvrier et al., 2022; Jonzén et al., 2010; Marescot et al., 2018), either due to abiotic natural environmental variability, biotic inter- and intraspecific interactions or due to anthropogenic disturbances (e.g. climate change or poaching). For example, the survival of cubs and subadults of spotted hyenas (*Crocuta crocuta*) in the Serengeti National Park declined strongly in response to a canine distemper virus outbreak in 1993/1994 (Benhaiem et al., 2018). Hunting, by targeting specific age classes, may disrupt the demographic structures of populations, leading to transient dynamics (Koons et al., 2005). For instance, the survival of black rhinoceros *Diceros bicornis* in north-West Namibia has increased after protection from illegal poaching (Brodie et al., 2011). As vital rates vary in time, the resulting demographic resilience is also expected to vary in time. However, the extent of such variation, when to account for it and the conditions that cause extreme fluctuations over time remain unknown. In a world where disturbances are on the rise (Turner, 2010), gaining a better understanding of the variations in the short-term responses to these disturbances can be crucial for conservation, wildlife management (Gerber & Kendall, 2016).

Here we propose the R package demres, which provides functions to quantify time-varying and time-constant demographic resilience, as well as to compare these two approaches both visually and by means of several distance metrics. We expect that demres will motivate quantification of time-varying demographic resilience for populations where relevant data is available. This, in turn, will facilitate comparative cross-study analyses that would ultimately assess for what species or populations and under what environmental conditions the assumption of time-constant demographic resilience holds.

### How to assess demographic resilience

Analyses of demographic resilience come from the framework of transient analyses (Caswell, 2006; Stott et al., 2011). This approach relies on (i) a matrix denoted **A** summarising the vital rates of the population and (ii) an initial demographic distribution representing the initial abundance distribution of each age or stage class of the study population; hereafter referred to as “(st)age” class for short. These two elements are thus key inputs that the user must provide to demres. Finally, using these two elements, the population is projected and its transient behaviour is quantified before it reaches asymptotic growth. These transient analyses allow calculating a set of demographic resilience metrics including convergence time, damping ratio, inertia, maximum amplification, maximum attenuation and reactivity (Table 1).

**Table 1:**
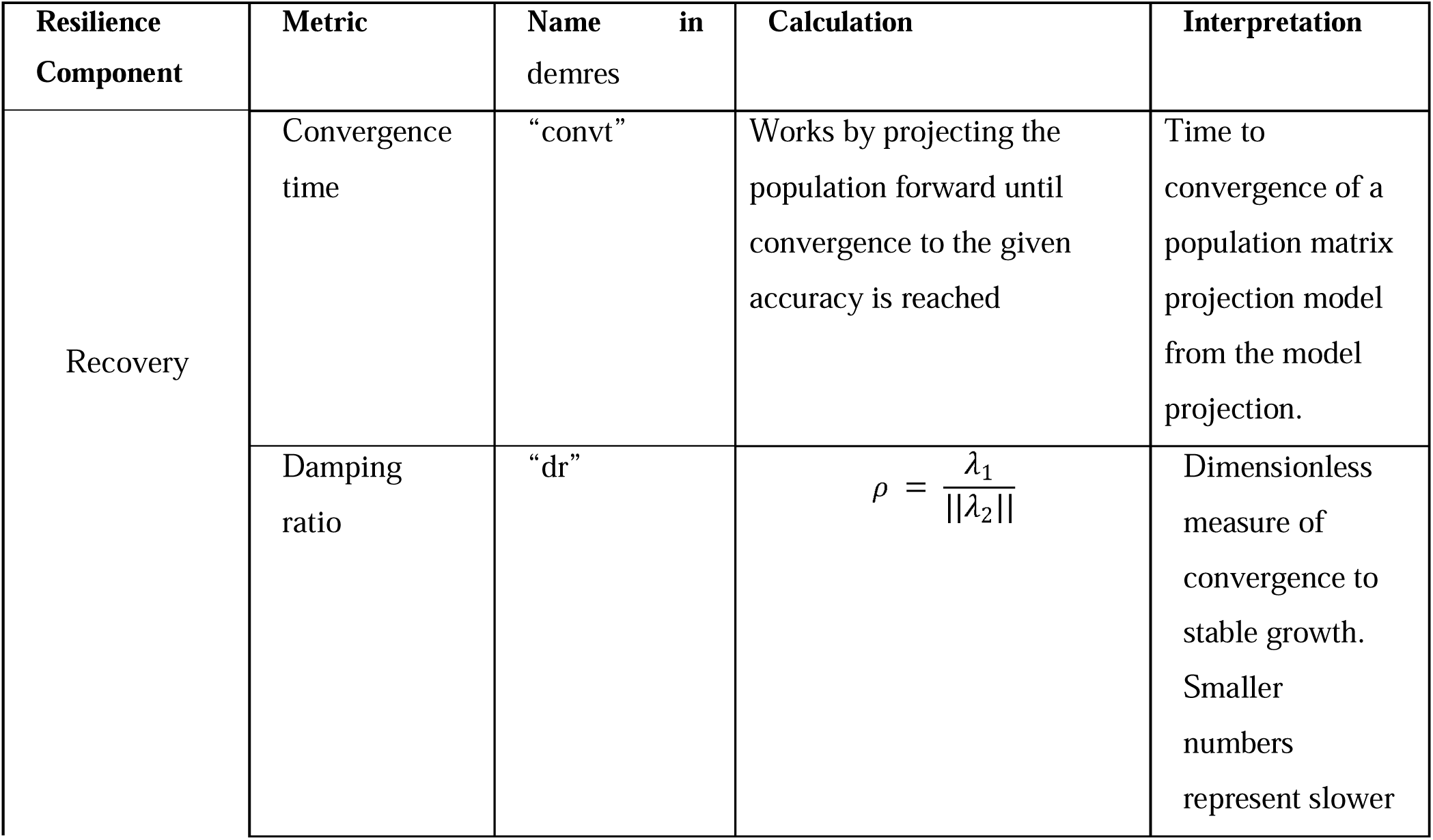
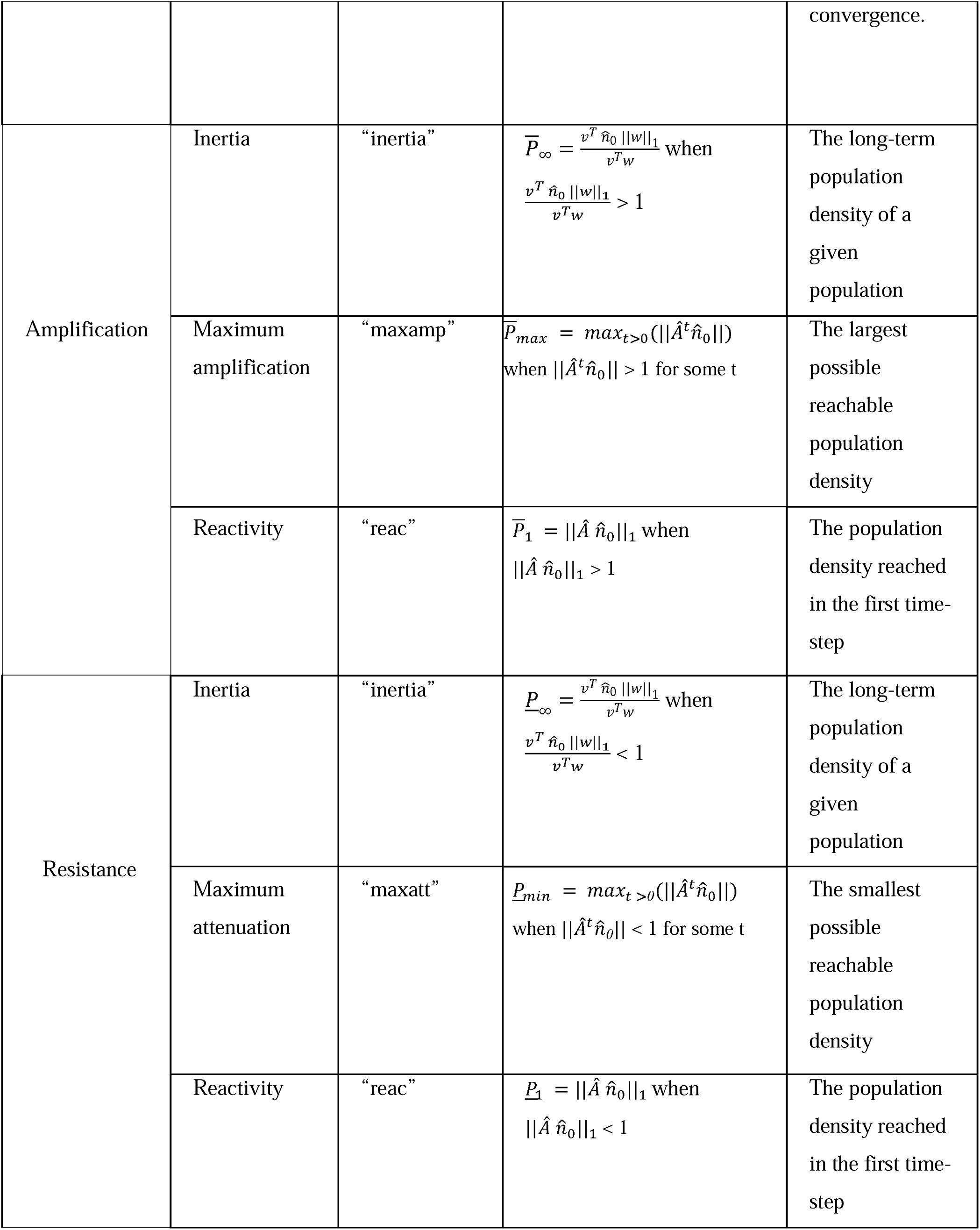
resilience metrics provided by demres, their calculation and interpretation. **A** is the population matrix. **Â** is the standardised matrix, which is calculated as **A**/,1_max_, where,1_max_ is the dominant eigenvalue of **A**. *w* is the dominant right eigenvector and the stable demographic structure of **A**. *v* represents the dominant left eigenvector, the reproductive value vector of **A**. The vector n’_0_ represents the initial demographic distribution, standardised to sum to 1.||m||_1_ is the one-norm of a vector *m* (equal to the sum of the entries for *m*). Following Stott et al. 2011, the case-specific demographic resilience metrics (i.e. calculated with a specific initial demographic distribution n’_O_) are represented with the Latin P (See appendix 1 for a similar table done for the bounds). An overbar (-) indicates an index of amplification, whereas an underbar (_) represents an index of attenuation. Transient metric subscripts provide information regarding the timeframe of a study, where 1 indicates first time-step; *max* and *min* are maximum amplification and attenuation, respectively, and ∞ is inertia.,1_1_ is the dominant eigenvalue,,1_2_ is the largest subdominant eigenvalue. Formulas, descriptions and table structure are based on Table 1 of Capdevila et al. (2020) and Stott et al. (2011). Interpretation of all metrics except convergence time and damping ratio are to be made relative to a population with stable growth rate.

### Matrix Population Models

In the fields of ecology and conservation biology, MPMs have become the most widespread tool to forecast population dynamics. The MPM is a mathematical representation of the change of distribution of individuals across different stages (e.g. size, or developmental stage) or age classes in a population over time (Caswell, 2006). The MPMs are built around a matrix **A** where the elements (known as vital rates) define, per unit of time, survival rates in a given (st)age, transition probabilities from one (st)age to another, and fecundity (i.e. per-capita number of offspring that is contributed to the population by each (st)age).

### Demographic distribution

The initial demographic distribution represents the abundance of individuals in each (st)age class by a vector (n’_0_). Such demographic distributions can be obtained from: i) a known (st)age-class distribution that is representative of the population being studied; or ii) stage-biased vectors that represent the most extreme cases where only one (st)age class is represented in the population, whereas abundances of all others are set to “0” (Townley & Hodgson, 2008).

Since demographic distribution often is not available for wild-living species, most existing studies apply option ii) to study resilience. Using stage-biased vectors provides the extreme plausible population responses to a disturbance, which are called the transient bounds (Stott et al., 2011; Townley & Hodgson, 2008). These bounds delineate the range of responses that a population can exhibit. Since stage-biased vectors are biased towards one (st)age class, there are as many stage-biased vectors as there are classes. For example, a population with four (st)age classes will have the four following standardised stage-biased vectors: [1 0 0 0]; [0 1 0 0]; [0 0 1 0]; [0 0 0 1]. The stage-biased vectors are automatically computed by demres, thus facilitating the transient analyses, especially if many (st)ages are distinguished.

### Standardising MPMs and vectors

Transient analyses can be performed in two ways (Stott et al., 2011). The first option uses absolute change in population abundance and thus describes the combined influence of both transient and asymptotic dynamics. The second option uses relative measures of transient dynamics, which allows disentangling the transient and asymptotic effects and therefore enables the comparison of populations with very different ranges of transient dynamics (Stott et al., 2011). For this, the matrix **A** is divided by,1_max_, the asymptotic growth rate, and the initial vector is divided by the total sum of individuals to get a total population density of 1. The package demres uses the matrices that are divided by,1_max_, but the users can decide to provide absolute or relative vectors.

### Comparison of the time-varying and the time-constant approaches

When data is collected from multiple years, or other biologically relevant time-steps, vital rates are calculated and matrices can therefore be built for each of these time-steps (Fig. 1A). Time-step for MPMs should be chosen so as to best reflect the life cycle of a study species (Enright et al., 1995). The time-varying approach considers vital rates calculated at each time-step (Fig. 1A). A matrix summarising these vital rates is then constructed for each of these time-steps and the transient analysis is performed on each of these matrices (Fig. 1B), providing resilience metrics at each time-step (Fig. 1D). In contrast, the time-constant approach relies on a matrix that is averaged over time (Fig. 1C) and thus returns only one value for each demographic resilience metric over the whole study period (Fig. 1E). For example, if a population was monitored for ten years and a time-step was a year, each of the demographic resilience metrics will be computed ten times following the time-varying approach, and once following the time-constant approach. Importantly, the average of the resilience metrics obtained with the time-varying approach is usually not equal to the resilience metric derived from the averaged population matrix (i.e. time-constant approach). The package demres allows for a visual comparison of the metric values obtained with the time-constant and time-varying approaches (Fig. 1F). To assess to what extent the demographic resilience metrics vary over time, demres computes the distance between the single value returned for the whole time series by the time-constant approach and all the time-specific values from the time-varying approach (Fig. 1G). Three distances can be computed: residual mean squared error (RMSE), relative residual mean squared error (rRMSE) and the mean absolute proportional error (MAPE, see demres vignette for formulas) (Hodson, 2022).

**Figure 1:**
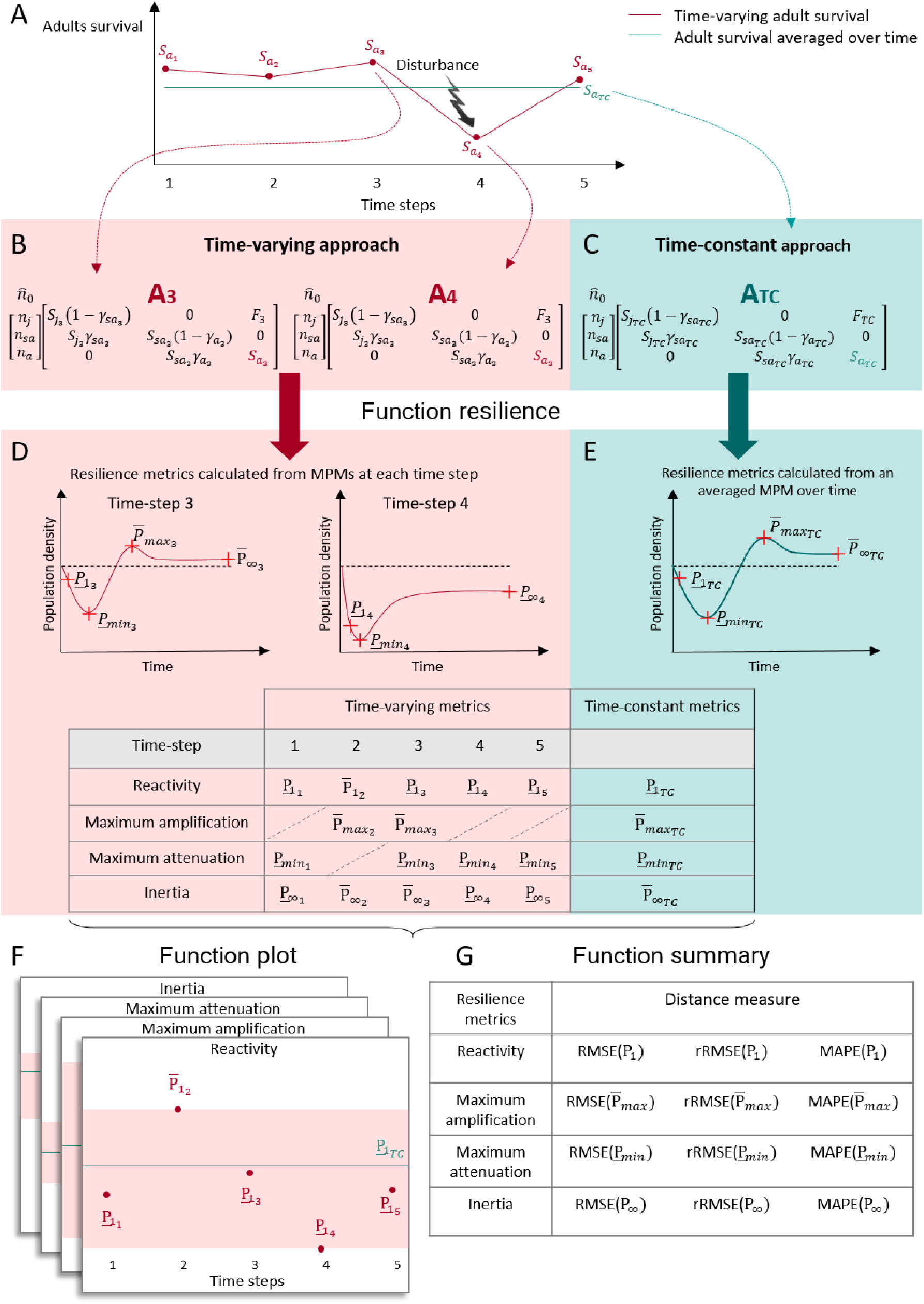
The workflow of the demres package. (A) In a hypothetical population with three stage classes; juveniles, subadults and adults, the vital rates vary over time. Here we display the survival of adults (S_a_), which is negatively affected by the disturbance that happens in the fourth year of the study. (B) The time-varying approach considers a matrix A_x_ for each time- step *x* and an initial demographic distribution n’_0_ and calculates the demographic resilience metrics at each time-step, capturing the temporal variation in vital rates. The index *x* indicates the time-step. The elements of the matrix A_x_ are: S_JX_, S_saX_, and S_aX_ - survival of juveniles, subadults and adults, respectively; and F_x_ - fecundity. y_saX_ and y_aX_ are the transition probabilities from juvenile to subadult and from subadult to adult, respectively. In n’_O_, n_J_, n_sa_ and n_a_ represent the relative proportion of juveniles, subadults and adults in the population. (C) The time-constant approach uses an averaged matrix A_TC_ that is obtained by averaging over the annual vital rate values. The index *TC* indicates the time-constant approach. (D) Given the initial population distribution and the matrix A_x_, the demographic resilience metrics are calculated, e.g. reactivity, maximum attenuation and inertia (see Table 1 for formulas). Following Stott et al. 2011, we used the Latin P to represent demographic resilience metrics calculated with a specific initial demographic distribution n’_0_ (in opposition to the Greek p used to represent the bounds, see appendix 1 for the formulas of the bounds). An overbar (-) indicates an index of amplification, whereas an underbar (_) represents an index of attenuation. The direction of the metrics have been chosen for illustration purposes only in this example. It is possible that the population does not amplify, as is the case here for the time-steps one, four and five. The population also did not attenuate when using the matrix LJ_2_ for the time-step two. (E) Under the time-constant approach the population attenuated and amplified. The package demres also provides: (F) a plot function visualising the resulting demographic resilience metrics, and (G) a summary function providing measures of distance between the time-varying and the time-constant metrics. The symbols shown in orange/blue denote, respectively, the metrics calculated under time-varying and time-constant approach.

### Resilience metrics

Demographic resilience is commonly quantified with a set of resilience metrics (Table 1, Capdevila et al. 2020, Stott et al. 2011). The main goal of demres is to compute them using either a time-varying or time-constant approach, or both.

### Package overview

The package demres will be available on GitHub: [true link will be added during proof] and will be submitted to CRAN. It is inspired and based on the popdemo package (Stott et al., 2012). The main function in demres is called resilience. We also programmed methods for the generic functions summary and plot working with the outputs produced by resilience. When designing demres we decided to follow the same syntax as in popdemo so as to facilitate an easy transition between the packages for the users.

### Calculate demographic resilience metrics based on a list of Matrix Population Models

The core function resilience allows calculating demographic resilience metrics using a list of matrices computed at each time-step of the study. The user can choose to compute a specific metric, several metrics, or all available metrics, including convergence time (“convt”), damping ratio (“dr”), inertia (“inertia”), maximum amplification (“maxamp”), maximum attenuation (“maxatt”) and reactivity (“reac”, Table 1). The choice of which metrics to use is done by setting the argument metrics. The user can choose to compute the metrics using either time-varying, time-constant or both approaches, using the argument time.

The initial demographic distribution can be specified in several ways, depending on the main goal of the analyses and the data availability. If the focus is on dynamics of a population with a particular demographic distribution (e.g. in reintroduction scenarios), then (i) this distribution can be supplied as one vector in vector (Fig. 2 A). If the goal is to assess resilience of the studied population over a certain time period in the past and demographic distribution for each time-step is available, the user can (ii) supply a list of vectors of (st)age- class distributions for each time-step in vector (Fig. 2 B). In cases when the interest lies in comparing resilience metrics across studies, the user can (iii) request bounds to be calculated from the (st)age-biased vectors by using bounds = TRUE or supplying manually one stage- biased vector in vector (Fig. 2 C). Finally, if the goal is, similarly to the option (ii) to assess resilience of a population in the past but not enough detailed data is available to extract demographic distribution for each time-step, then (iv) a population can be projected from a given initial demographic distribution and a list of matrices over a certain time period, using TDvector = TRUE for Time-Dependent vector. The demographic distribution for the following time step is then projected based on the provided initial demographic distribution and the respective matrix for that time-step. This procedure reiterates across all time-steps (Fig. 2 D).

**Figure 2:**
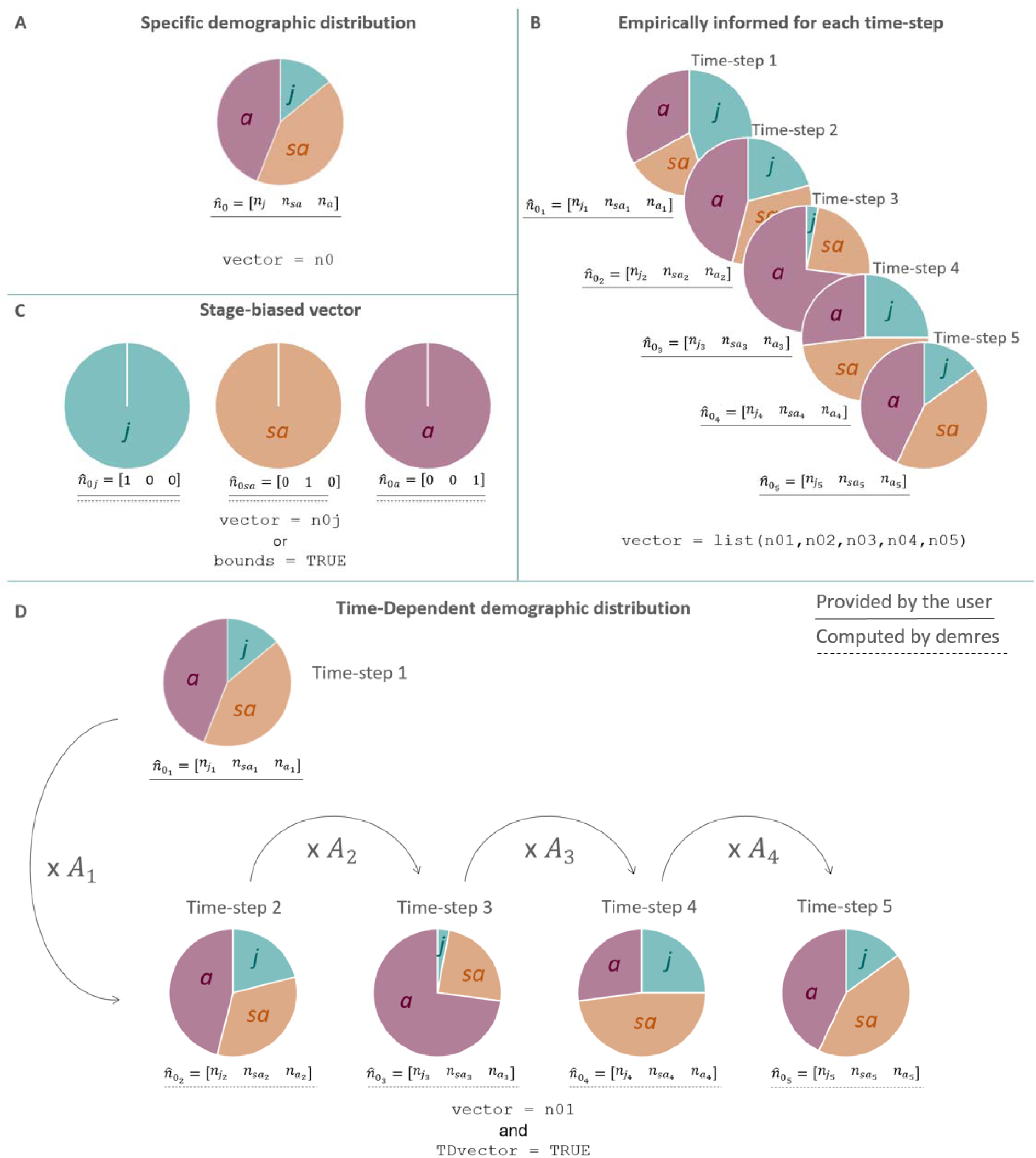
Representation of the different options a user has when specifying the demographic distribution when studying a population structured in three stage classes juveniles, subadult and adults. (A) In the case of assessing the demographic resilience of a population with a specific abundance of juveniles (represented as *j*), subadults (*sa*) and adults (*a*), then the user can provide in vector with,, the relative density of juveniles, subadults and adults in the population, respectively. (B) In the case where the demographic distribution is known for each time-step, then the user can provide a list of vectors n·_0_ for each time-step *x* in vector. (C) In the case of assessing the transient bounds, the user can either provide the stage-biased vector and test for each one of them or use the argument bounds specified to return the bounds of each of the resilience metrics. Finally, if a specific demographic structure is known but then needs to be projected over each time-step, then the user can provide n’_O_ in vector for the first time-step and use the argument TDvector for demres to compute the initial vectors for the rest of the time-steps.

## Case study

### Description of the blue crane study

The data distributed with this package (bluecrane) comes from a study by (Altwegg & Anderson, 2009) of a blue crane population (*Anthropoides paradiseus*) in South Africa. This population was monitored between 1997 and 2008, with a total of 451 individuals ringed to assess how rainfall impacted vital rates. Five age classes were distinguished. The study revealed that the survival of blue cranes in all age classes increased with increasing rainfall in the late breeding season, and varied substantially over the 12 years of the study, with lowest values in the fifth year and highest values in the second year of the study. The reproductive output of blue cranes also varied substantially over time, being higher in the years with higher rainfall during the early breeding season. As a result, we expected demographic resilience to vary with time.

### Assess demographic resilience metrics

We extracted the 12 MPMs for each time-step of the study from the COMADRE database (SalgueroLJGómez et al., 2016). COMADRE contains MPMs of animal species, which can be directly downloaded from their website (https://compadre-db.org/) or using the R package Rcompadre (Jones et al., 2022).

For illustration purposes, we here focus on quantifying a single resilience metric: reactivity. Since this species is vulnerable (IUCN 2024) and reproduction can be lowered due to climate, we use an initial demographic distribution with an under-representation of the first age class. The function resilience is used to specify the list of matrices with the argument listA and the metric to be calculated (“reac”) with the argument metrics. If the user wants to compute the bounds, then the argument bounds should be set to TRUE. Finally, if the metric is to be calculated using both time-varying and time-constant approaches then time should be set to “both”:

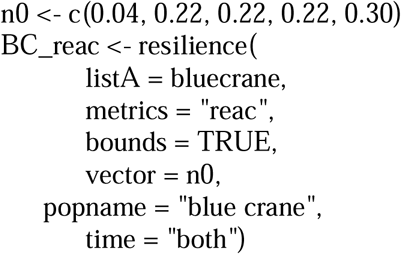

The time-constant reactivity obtained is 1.31 (Table 2) with the lower bound at a value of 0.53 and the upper bound at a value of 2.90. A reactivity of 1.31 means that as the immediate response to a disturbance the population will grow 1.31 times faster than its stable growth rate. The time-varying reactivity varies between 1.22 (time-step 5) and 1.44 (time-step 2) when calculated with the supplied initial vector. The lower bound varies between 0.49 (time- step 5) and 0.57 (time-step 2) while the upper bound varies between 2.52 (time-step 5) and 3.38 (time-step 2). The time-varying reactivity differs most from the time-constant value when using the upper bound (Fig. 3). This conclusion is also supported by RMSE and MAPE, whose values are largest when calculated at the upper bound (Table 3).

**Figure 3:**
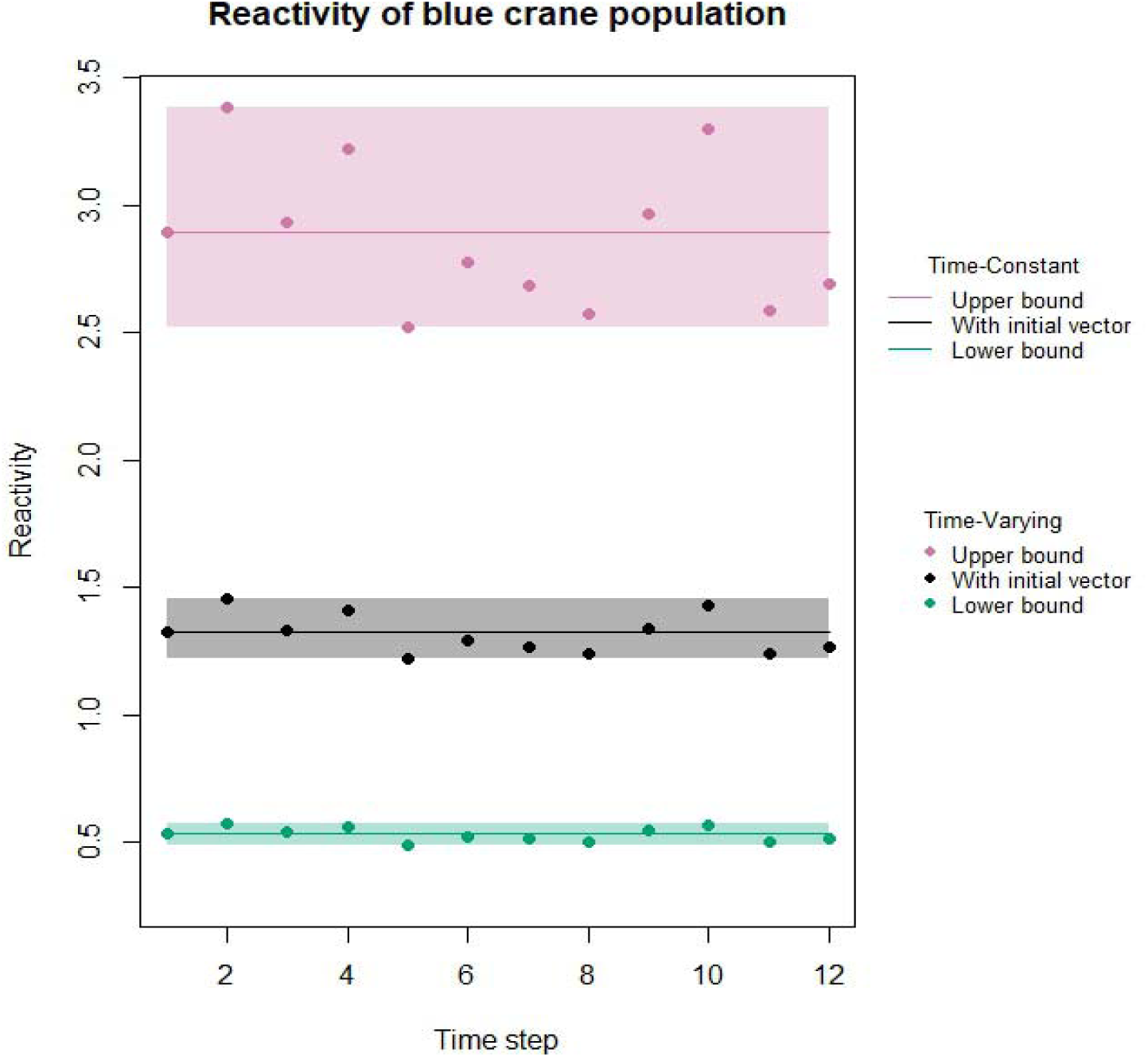
Application of the function plot to visualise time-varying (dots) and time-constant (solid lines) reactivity values of the blue crane population. The reactivity is calculated with the specified initial vector and its lower and upper bounds. The shaded blocks represent the range between the minimum and the maximum of the time-varying values.

**Table 2:** Reactivity of the blue crane population calculated using both approaches (time- varying: TV, time-constant: TC). Values that were calculated using the initial demographic distribution are shown as reac, and those calculated using the (st)age-biased vectors are shown as: reac_lwr and reac_upr (for lower and upper bound, respectively).This table is the output produced by the function resilience.

| Time-step | Population name | reac_TV | reac_lwr_TV | reac_upr_TV | reac_TC | reac_lwr_TC | reac_upr_TC |
| --- | --- | --- | --- | --- | --- | --- | --- |
| 1 | blue crane | 1.31 | 0.53 | 2.89 | 1.31 | 0.53 | 2.89 |
| 2 | blue crane | 1.44 | 0.57 | 3.38 | 1.31 | 0.53 | 2.89 |
| 3 | blue crane | 1.32 | 0.54 | 2.93 | 1.31 | 0.53 | 2.89 |
| 4 | blue crane | 1.40 | 0.56 | 3.22 | 1.31 | 0.53 | 2.89 |
| 5 | blue crane | 1.22 | 0.49 | 2.52 | 1.31 | 0.53 | 2.89 |
| 6 | blue crane | 1.28 | 0.52 | 2.78 | 1.31 | 0.53 | 2.89 |
| 7 | blue crane | 1.26 | 0.52 | 2.68 | 1.31 | 0.53 | 2.89 |
| 8 | blue crane | 1.23 | 0.50 | 2.58 | 1.31 | 0.53 | 2.89 |
| 9 | blue crane | 1.33 | 0.55 | 2.96 | 1.31 | 0.53 | 2.89 |
| 10 | blue crane | 1.42 | 0.57 | 3.30 | 1.31 | 0.53 | 2.89 |
| 11 | blue crane | 1.23 | 0.50 | 2.59 | 1.31 | 0.53 | 2.89 |
| 12 | blue crane | 1.26 | 0.51 | 2.69 | 1.31 | 0.53 | 2.89 |

**Table 3:**
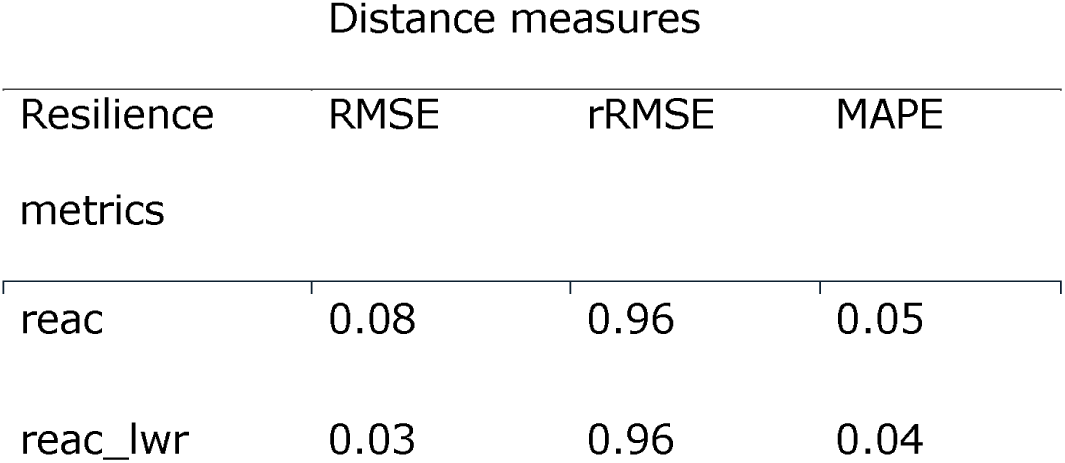
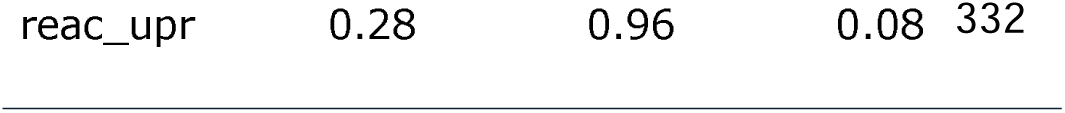
The distance measures between the time-varying and the time-constant values of reactivity calculated for the blue crane population. This table was obtained by calling the function summary on an object returned by the function resilience.

One application of the demographic resilience approach for conservation is to investigate which (st)age class affects the resilience metrics the most. For the blue crane population the upper bound of reactivity is reached when only the oldest individuals (“year 5 and older”) are present (see vignette). This implies that this population will grow faster than the asymptotic rate (i.e. amplify) when there is an overrepresentation of such individuals. Identifying the specific (st)age classes and vital rates that can have profound effects on variation in resilience of the overall population is of great importance for conservation. In the case of the blue crane, our findings suggest that a population reinforcement aiming at increasing the stock of “older” individuals could be used to counter the negative effects of climate change.

## Conclusions

The demres package provides tools to both assess time-varying demographic resilience metrics and to compare time-varying and time-constant approaches visually and quantitatively. Our package draws attention to the time-varying character of resilience, and permits pinpointing time intervals over which the values of demographic resilience metrics were extreme. We provide a flexible piece of software that can be easily used by conservation biologists, population ecologists, and managers who work on long-term studies and aim at assessing the resilience of the studied population. The framework we provide allows comparing demographic resilience metrics in a standardised way, facilitating the comparison between populations or species in comparative studies.

## Code availability

The R package demres is freely available on GitHub: https://github.com/JulieLouvrier/demres. We also intend to submit it to CRAN.

## Data availability

The data attached with the package can be found in the package repository in Github, or by calling data(bluecrane) after loading the package.

## Conflict of interest

The authors declare no conflict of interest.

### Acknowledgements

We thank Oliver Höner from the WILDER project for his critical comments on the manuscript., JL, EWW, ATC, SB and VR acknowledge BMBF project WILDER (Number 16DKWN148) for financial support.

## Authors contributions

JL conceptualization, package development, writing, AC package development, writing, EWW package development, writing, ATC conceptualization, writing, SB conceptualization, writing, VR conceptualization, package development, writing.

## Notes

### Competing Interest Statement

The authors have declared no competing interest.

