## Appendix 1 for "demres: An R package to study time-varying demographic resilience"

| **Resilience Component** | **Metric** | **Name in** demres | **Calculation** | **Interpretation** |
| --- | --- | --- | --- | --- |
| Recovery | Convergence time | “convt_upr”  &  “convt_lwr” | Works by projecting the population forward until convergence to the given accuracy is reached | Minimum and maximum time to convergence of a population matrix projection model from the model projection obtained with the stage-biased vectors. |
|  | Damping ratio | “dr” | $\rho= \frac{\lambda_{1}}{\vert\vert\lambda_{2}\vert\vert}$ | Dimensionless measure of convergence to stable growth. Smaller numbers represent slower convergence. |
|  | Inertia | “inertia_upr” | $\bar{\rho}_{\infty}= \frac{{v_{max}\vert\vert w\vert\vert}_{1}}{v^{T}w}$ | The largest possible long term population density. |
| Amplification | Maximum amplification | “maxamp_upr” | $\bar{\rho}_{max}=\frac{max}{t>0}({\vert\vert\hat{A}^{t}\vert\vert}_{1})$ | The largest population density that can be reached at any time after disturbance. |
|  | Upper Reactivity | “reac_upr” | $\overline{\rho}_{1}={\vert\vert\hat{A}\vert\vert}_{1}$ | The largest population density that can be reached in the first time step after disturbance. |
| Resistance | Inertia | “inertia_lwr” | $\underline{\rho}_{\infty}=\frac{{v_{min}\vert\vert w\vert\vert}_{1}}{v^{T}w}$ | The lowest possible long term population density. |
|  | Maximum attenuation | “maxatt_lwr” | $\underline{\rho}_{min}= \frac{min}{t>0} (minCS(\hat{A}^{t}) )$ | The lowest population density that can be reached at any time after disturbance. |
|  | Reactivity | “reac_lwr” | $\underline{\rho}_{1}= minCS(\hat{A})$ | The lowest population density that can be reached in the first time step after disturbance. |

A is the matrix population model. Â is the standardized matrix population model, which is calculated as A/$\lambda_{max}$, where $\lambda_{max}$ is the dominant eigenvalue of A. *w* is the dominant right eigenvector and the stable demographic structure of A. *v* represents the dominant left eigenvector, the reproductive value vector of A. The vector $\hat{n}_{0}$represents the initial demographic distribution, standardized to sum to 1. minCS denotes the minimum column sum of a matrix, and ${||m||}_{1}$ is the one-norm of a vector m (equal to the sum of its entries). An overbar (¯) or underbar (_)indicate amplification and attenuation, respectively. Subscripts provide information regarding the timeframe of a study, where 1 indicates first time-step indices; max and min are maximal amplification or attenuation, respectively, and ∞ is inertia. λ1 is the dominant eigenvalue, λ2 is the largest subdominant eigenvalue.
